# Thermal energetics of bats of the family Vespertilionidae: an evolutionary approach

**DOI:** 10.1101/2024.01.03.574072

**Authors:** Jorge Ayala-Berdon, Kevin I. Medina-Bello, Jorge D. Carballo-Morales, Romeo A. Saldaña-Vázquez, Federico Villalobos

## Abstract

1. Thermal energetics define the way animals spend energy for thermoregulation. In this regard, numerous studies have determined that body mass (*M_b_*) is the most influential morphological trait affecting the thermal traits in different species of birds and mammals. However, most of the studies have been focused on the basal metabolic rate (*BMR*), while other thermal traits have been less studied.
2. We addressed this gap by examining thermal variables on bats of the family Vespertilionidae. Using open-flow respirometry, we measured *BMR*, absolute thermal conductance (*C*’), lower and upper critical temperatures (*T_LC_* and *T_UC_*), and the breadth of the thermoneutral zone (*TNZ_b_*) of 15 bat species varying in *M_b_* from ∼ 4.0 to 21.0 g from central Mexico. We: 1) combined our empirical data with information gathered from the bibliography and conducted phylogenetic analyses to investigate the relationship between *M_b_* and thermal traits, and 2) mapped the thermal energetic values along the phylogeny to explore how they may have evolved.
3. We found a positive relationship between *M_b_* and *BMR* and absolute *C’*, and a negative relationship between *M_b_*and *T_LC_* and *T_UC_*. However, we did not find a relationship between *M_b_*and *TNZ_b_* in bats. The phylogenetic approach suggested that over the evolutionary history of bats, *BMR* and *C*’ have decreased while *T_LC_* and *T_UC_*have increased.
4. Our results suggest that adaptive changes in *M_b_* and thermal traits may have influenced the geographical distribution and the use of energy-saving strategies of the different species of bats of the family Vespertilionidae.

## INTRODUCTION

Homeotherm endotherms have evolved the capacity to sustain their internal physiological processes optimally, regardless of changes in ambient temperature (*T_a_*) (Crompton et al., 1978). Nevertheless, endogenous homeothermy incurs important energetic costs associated with maintaining a high body temperature (*T_b_*) (Nagy et al., 1999; Speakman & Thomas, 2003). As the metabolic rate of individuals changes proportionally to the difference between *T_b_*and *T_a_* (Δ*T*°) for *T_b_* to remain constant (McNab, 2012) and *T_b_* is generally different from *T_a_* in the environment, a substantial proportion of animals’ energy is expended in thermoregulation (Speakman & Thomas, 2003; Geiser, 2021).

Endotherms have the capacity to modify their internal heat gain and loss, and consequently their metabolic rate, to some extent. To achieve this, animals change their thermal conductance (*C*’), adjusting their peripheral blood flow, flattening or erecting their fur or feathers, or modifying their posture to change the surface-area exposed to the environment (Speakman & Thomas, 2003). These modifications enable the individuals to present a constant metabolic rate even when *T_a_* changes (Heinrich, 1977; Young et al., 1989; McNab, 2012). The range of *T_a_*’s where the metabolic rate is constant can be measured in an adult animal at rest in a postabsorptive state under laboratory conditions. This range, known as the thermoneutral zone (*TNZ*), represents the minimum metabolic rate compatible with *T_b_* regulation (i.e., the basal metabolic rate -*BMR*-). Below and above the *TNZ*, the metabolic rate increases, as changes in isolation and posture no longer compensate for Δ*T*°. The *T_a_* values where animals start spending energy either to increase metabolic heat production at low *T_a_* values or to prevent overheating at high *T_a_* values are known as the lower critical temperature (*T_LC_*) and the upper critical temperature (*T_UC_*) (Hill et al., 2004; Withers et al., 2016). Both *T_LC_* and *T_UC_* define the breadth of the *TNZ* (*TNZ_b_*). Thus, the *TNZ_b_* represents the range of *T_a_* values where animals expend minimum energy for thermoregulation.

The thermal variables presented above (i.e., the *C*’, *BMR*, *T_LC_*, *T_UC_*, and *TNZ_b_*) can be used to characterize different species of endotherms. According to McNab (2012), this characterization is possible not because the individuals spend too much time in the resting and postabsorptive state, but because the conditions under which these variables are measured in the laboratory are the same for all endotherms. In this regard, many investigations have been conducted to measure thermal variables in different species of birds and mammals (Pollock et al., 2019; Andreasson et al., 2020; Pollock et al., 2021; Speakman & Thomas, 2003; Willis et al., 2005; Machado & Soriano, 2007; KsiaLzek et al., 2009, among others). These studies have shown that body mass (*M_b_*) is by far the most influential morphological trait affecting the *BMR* of individuals (McNab, 2008, McNab, 2009; Speakmand & Thomas, 2003). However, most of the research has been focused on the *BMR* of animals, while the other thermal variables have been less studied. In the last decades, Riek & Geiser (2013) evaluated the relationship between *M_b_* and *T_LC_*, *T_UC_*, and *TNZ_b_* of 203 mammal species. The authors found that all thermal variables were related to *M_b_*. Nevertheless, because the study was done on a high scale, this can mask the allometric relationships that can be observed in more specific groups of animals. Because thermal variables define the *T_a_* when individuals start spending energy (Reik & Geiser, 2013), their determination would help understand the relationship of the thermoregulatory traits of the species with their ecology, behavior, and geographical distribution (Gazier, 2010; Riek & Geiser, 2013; Bozinovic et al., 2014).

Bats is a well-suited group of mammals to study thermal energetics because of their diverse patterns of thermal regulation. Some species are obligate homeotherms while others can depress their metabolic rate below *BMR* to use daily torpor or hibernation (Stawski et al., 2014; Geiser, 2021). Among these, bats of the family Vespertilionidae account for one-third of the total species composing the order Chiroptera (Barclay & Harder, 2003). In these species, *M_b_* ranges from ∼ 2 to 90 g (Nowak & Walker, 1994; Moyers Arévalo et al., 2018). Therefore, their thermal variables should be related to *M_b_*. To evaluate this hypothesis, we measured the thermal energetics of 15 species of bats of the family Vespertilionidae, differing in *M_b_* from ∼ 4.0 to 21.0 g, living in central Mexico. Using our empirical data and information reported in the bibliography, we tested the relationship of *M_b_* and thermal traits of bats using a phylogenetic approach. We also modeled the thermal variables and mapped the traits along the phylogeny to explore how these traits may have evolved, and if they would be shared among closely related species. Because there is strong evidence that *M_b_* and some thermal traits like *BMR* have co-evolved in mammals (White et al., 2019), we predicted that thermal variables should be related to the evolution of the bats’ *M_b_*.

## MATERIALS AND METHODS

### Bat care and housing

Bats were captured from October to February between 2019 and 2022 at eight sites located in Central Mexico (Fig. 1) (Table 1). Mist nets (2 x 3, 2 x 6, and 2 x 9 m) were deployed near water bodies or within the vegetation at the study sites. Nets were opened at dusk and closed at ∼1:00 am. Non-reproductive adult males from 15 species of the Vespertilionidae family differing in *M_b_* were selected (Table 2). Some of these species were captured in small numbers due to their naturally low population size in the ecosystems (LaVal, 2004; Perry & Carter, 2010). Adult non-reproductive bats were chosen to avoid interference from the energy requirements associated with growth and reproduction (Genoud et al., 2018). Captured individuals were identified to species level using a bat identification guide for Mexico (Medellín et al., 2008). We assigned the taxonomic identities of the bat species following Ramírez-Pulido et al., (2014). Body mass (*M_b_*), length of the forearm, age (either juvenile or adult) and reproductive condition were recorded. Body mass was calculated using a digital balance to the nearest 0.2 g (Ohaus, Newark, NJ, USA). Forearm length was measured with a caliper to the nearest 0.1 mm. Age was determined by observing the epiphyseal space of the fourth metacarpal of the third and fifth fingers, since juvenile individuals present some visible space, or the space is not completely closed. Reproductive condition was determined by visualizing the testes, which in males increase in size during spermatogenesis (Wilkinson & Brunet-Rossinni, 2009).

**FIGURE 1.**
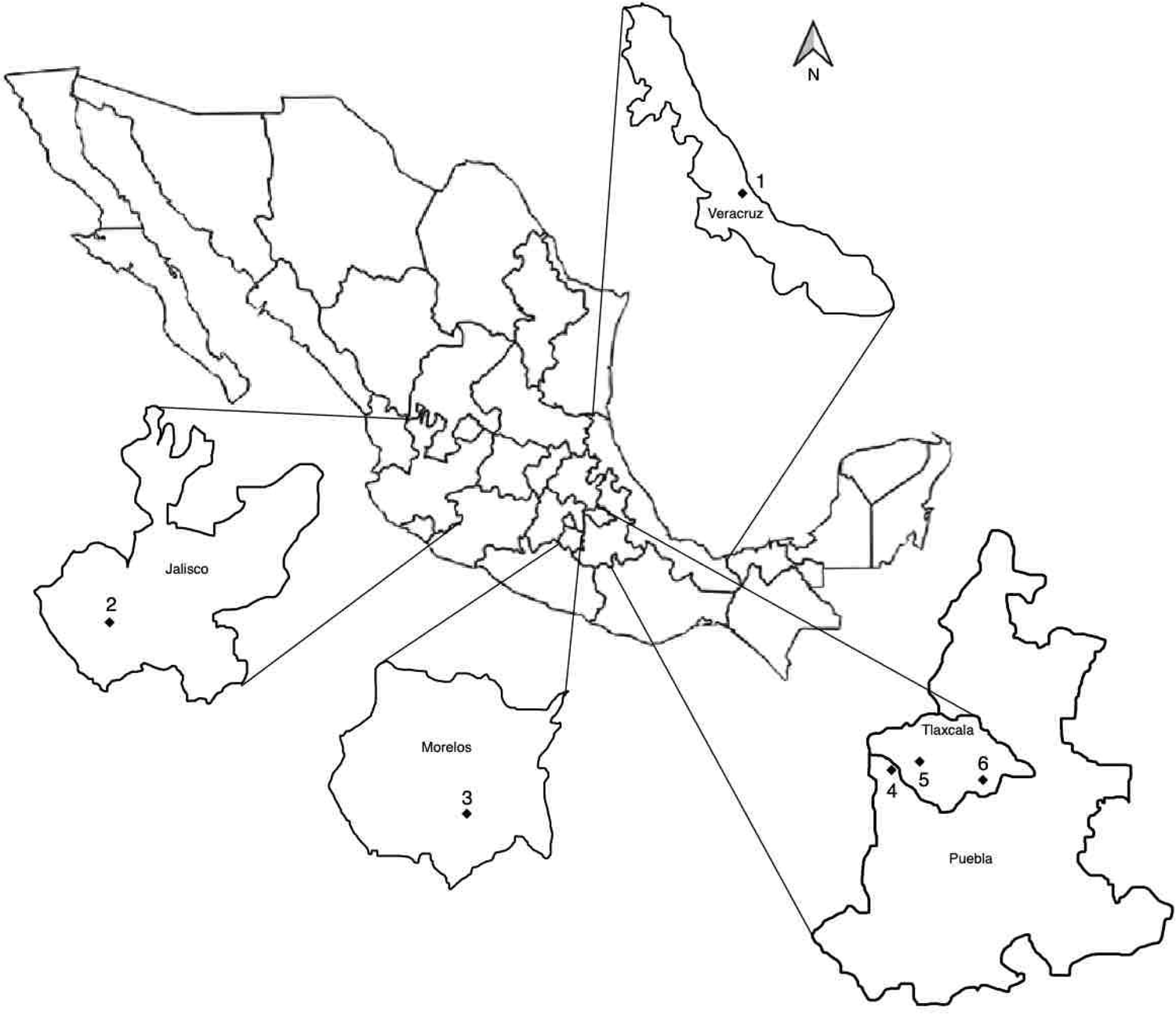
Bats were captured in six locations from five states in central Mexico. The main vegetation type, minimum average *T_a_*, and mean annual precipitation of localities are presented in table 1.

**TABLE 1.** Environmental characteristics of the study sites (Fig. 1) where bats of the family Vespertilionidae were captured for the measurements of thermal traits.

| Locality, state | Vegetation type | Mean annual<br>minimum $T_a$<br>(°C) | Mean annual<br>precipitation<br>(mm) | References |
| --- | --- | --- | --- | --- |
| La Mancha, Veracruz | Sub-deciduous forest | 19.0 | 744 | INEGI, 2005;<br>2017 |
| Autlán, Jalisco | Tropical dry forest | 13.6 | 861 | INEGI 2014;<br>2017 |
| SHBR, Morelos | Tropical dry forest | 15.5 | 665 | INEGI 2014;<br>2017; Chávez<br>2019 |
| SCM, Puebla | Coniferous forest | 8.0 | 1,400 | INEGI 2014;<br>2017 |
| Hueyotlipan, Tlaxcala | Sabino forest | 8.5 | 1,018 | INEGI, 2005;<br>2017 |
| LMNP, Tlaxcala | Coniferous forest | 5.0 | 1,150 | Villers et al.,<br>2006; INEGI<br>2005; 2017 |

SHBR= Sierra de Huautla Biosphere Reserve; SCM= Santa Cruz Moxolahuac; LMNP= La Malinche National Park

**TABLE 2.**
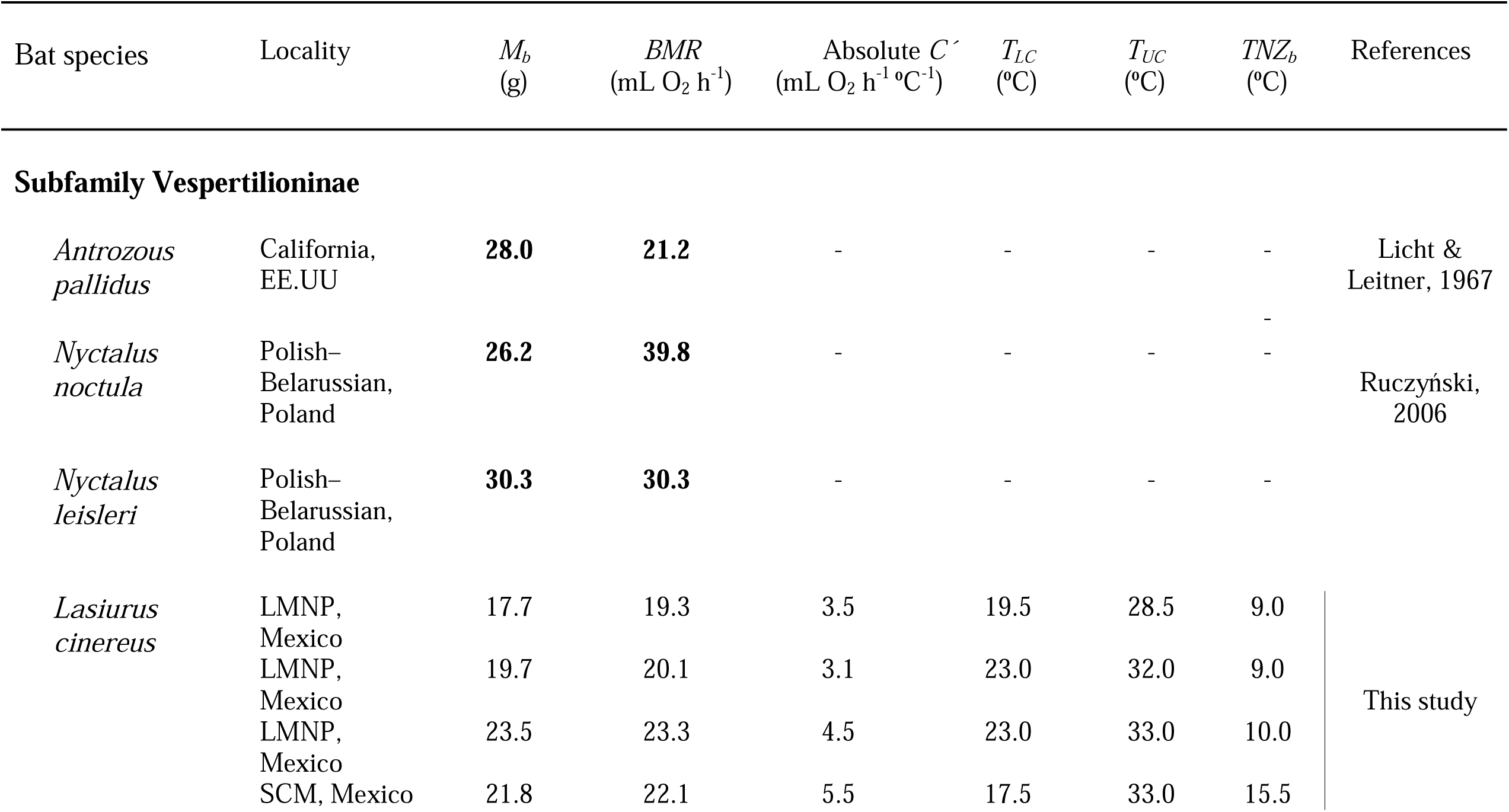
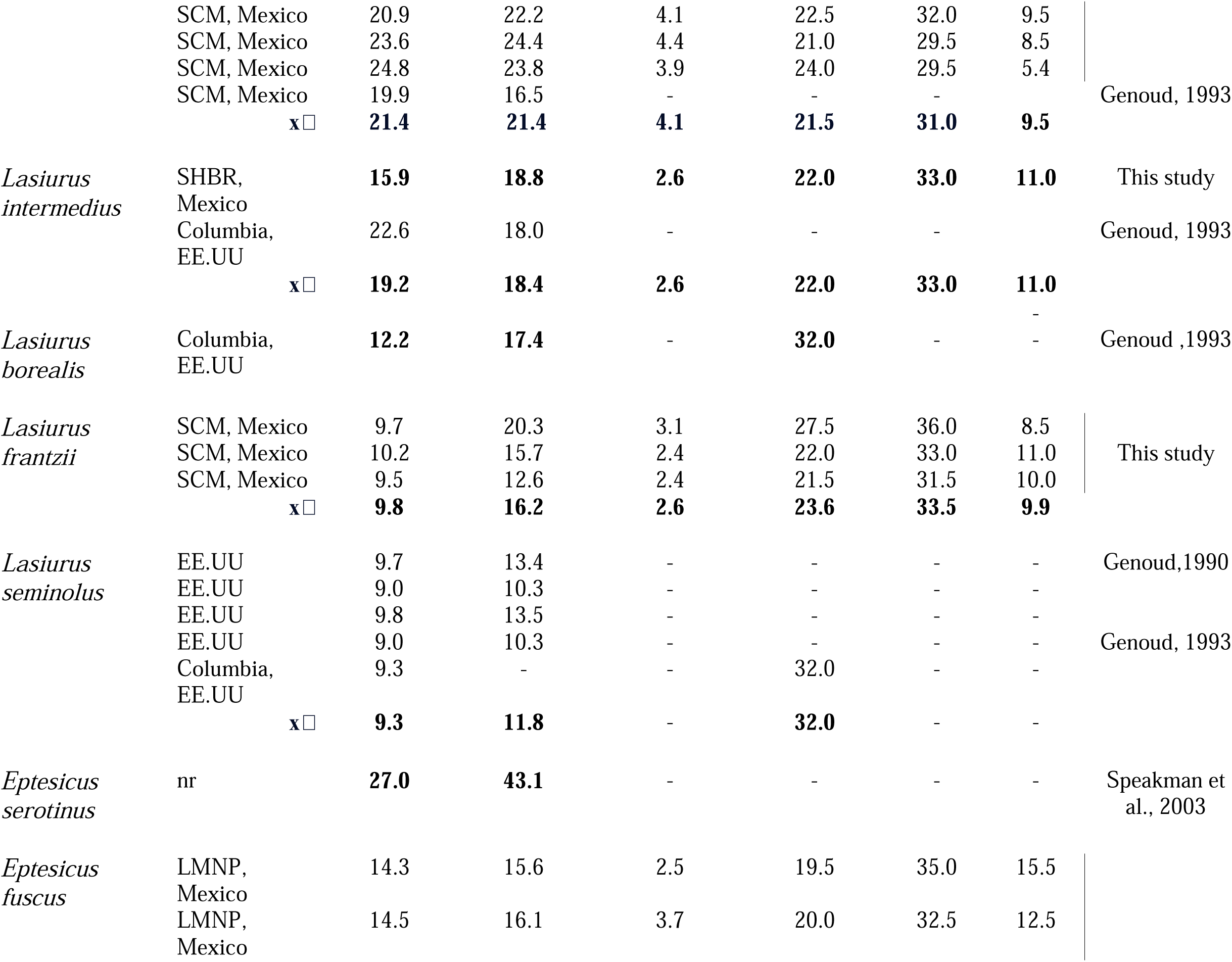
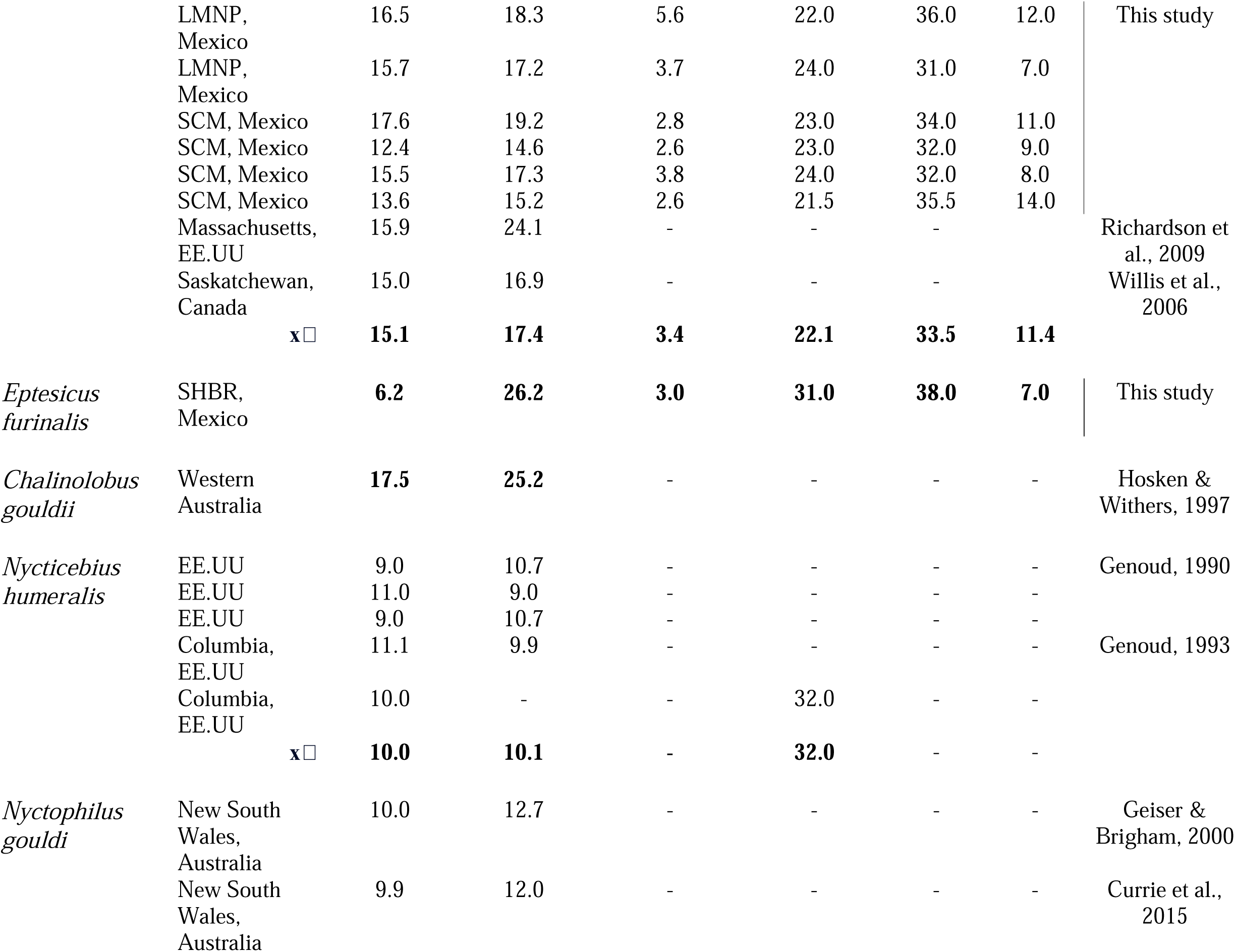
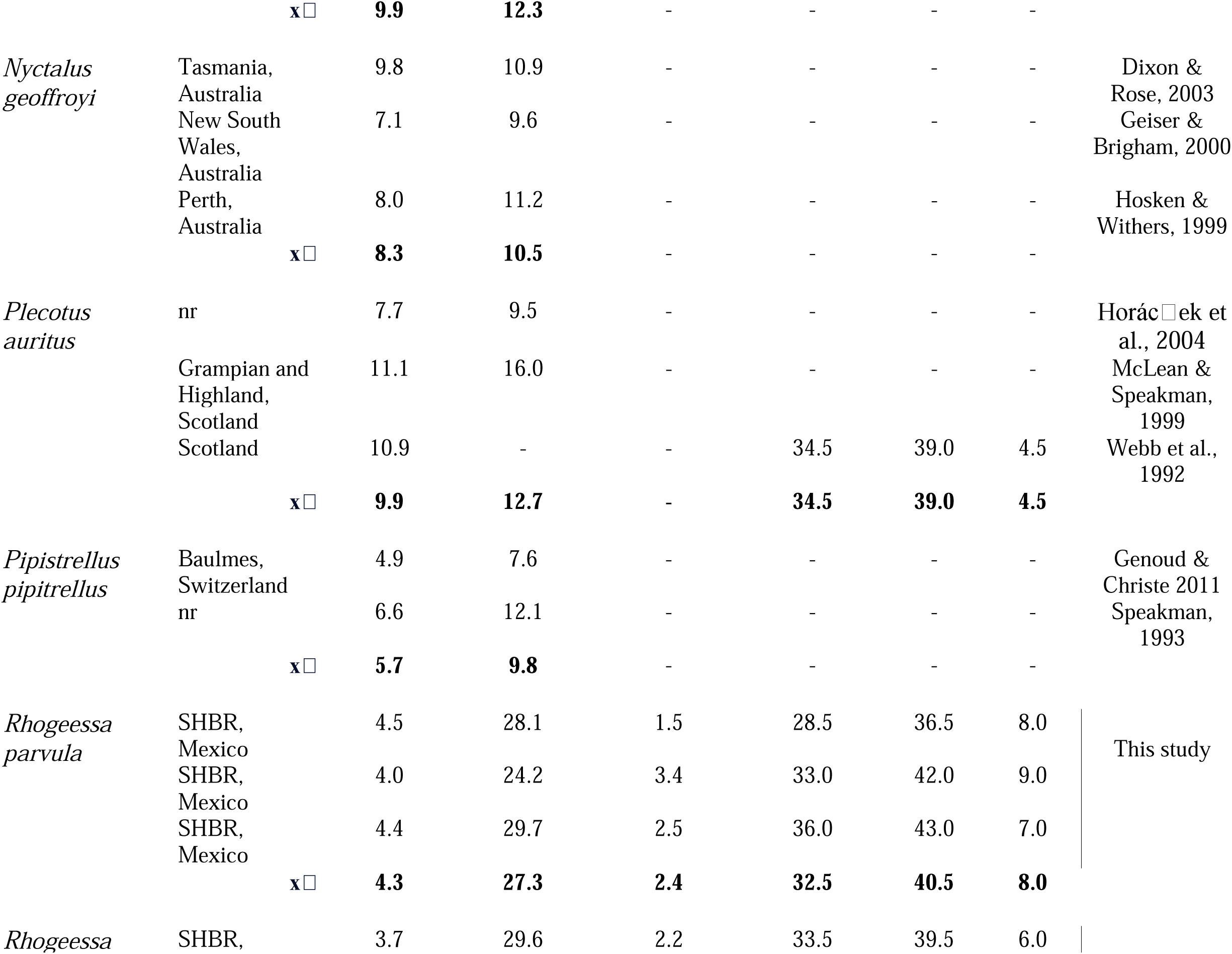
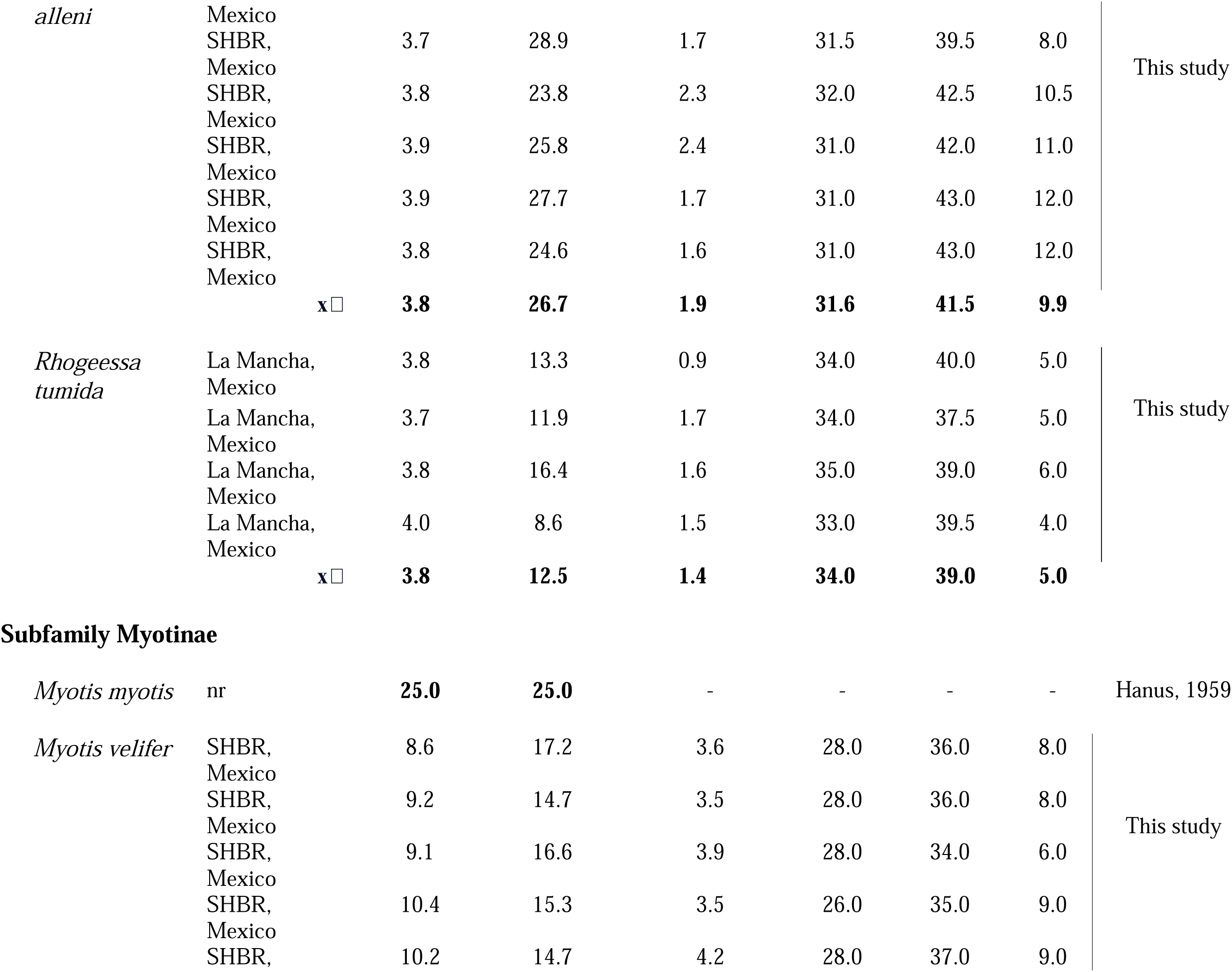
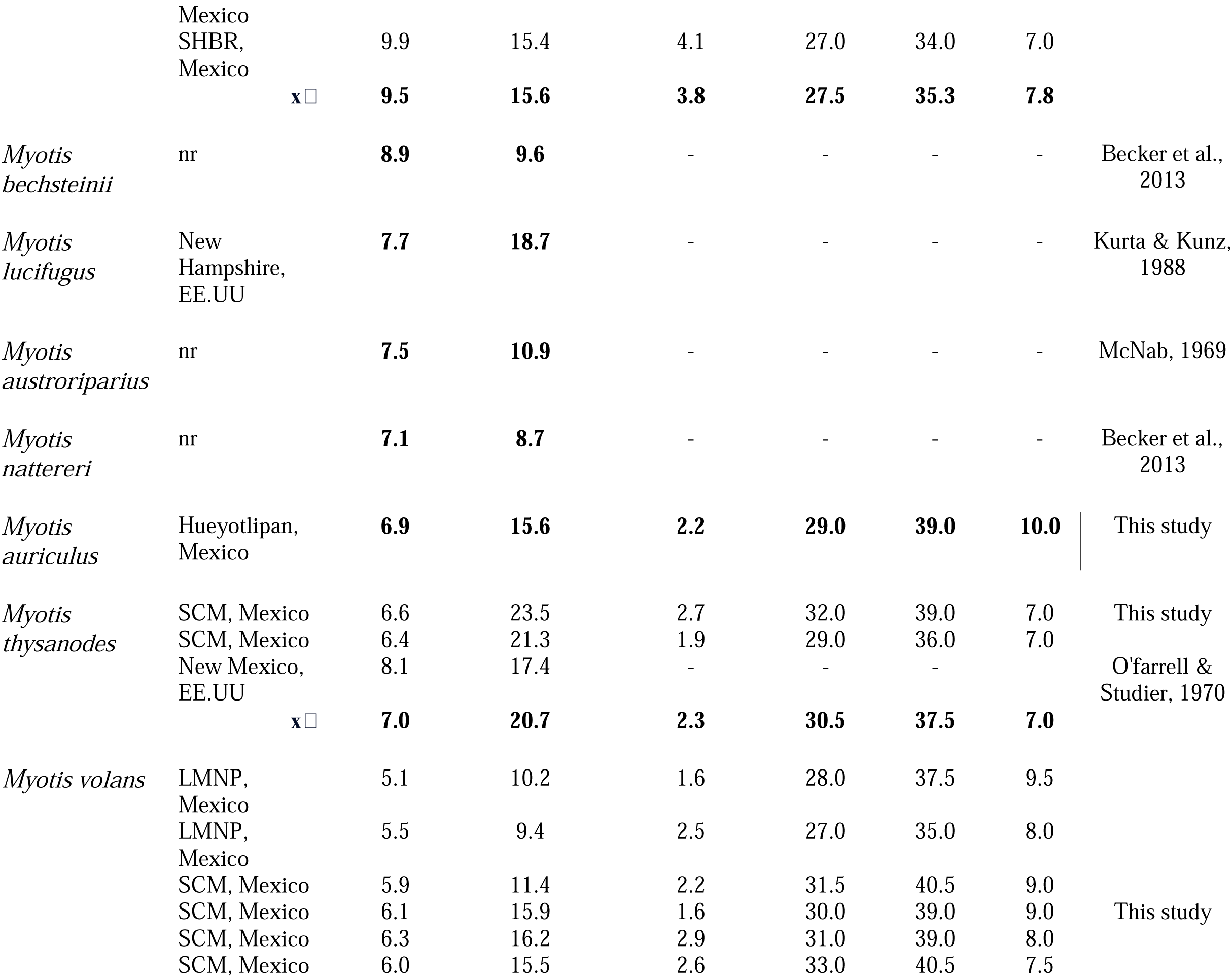
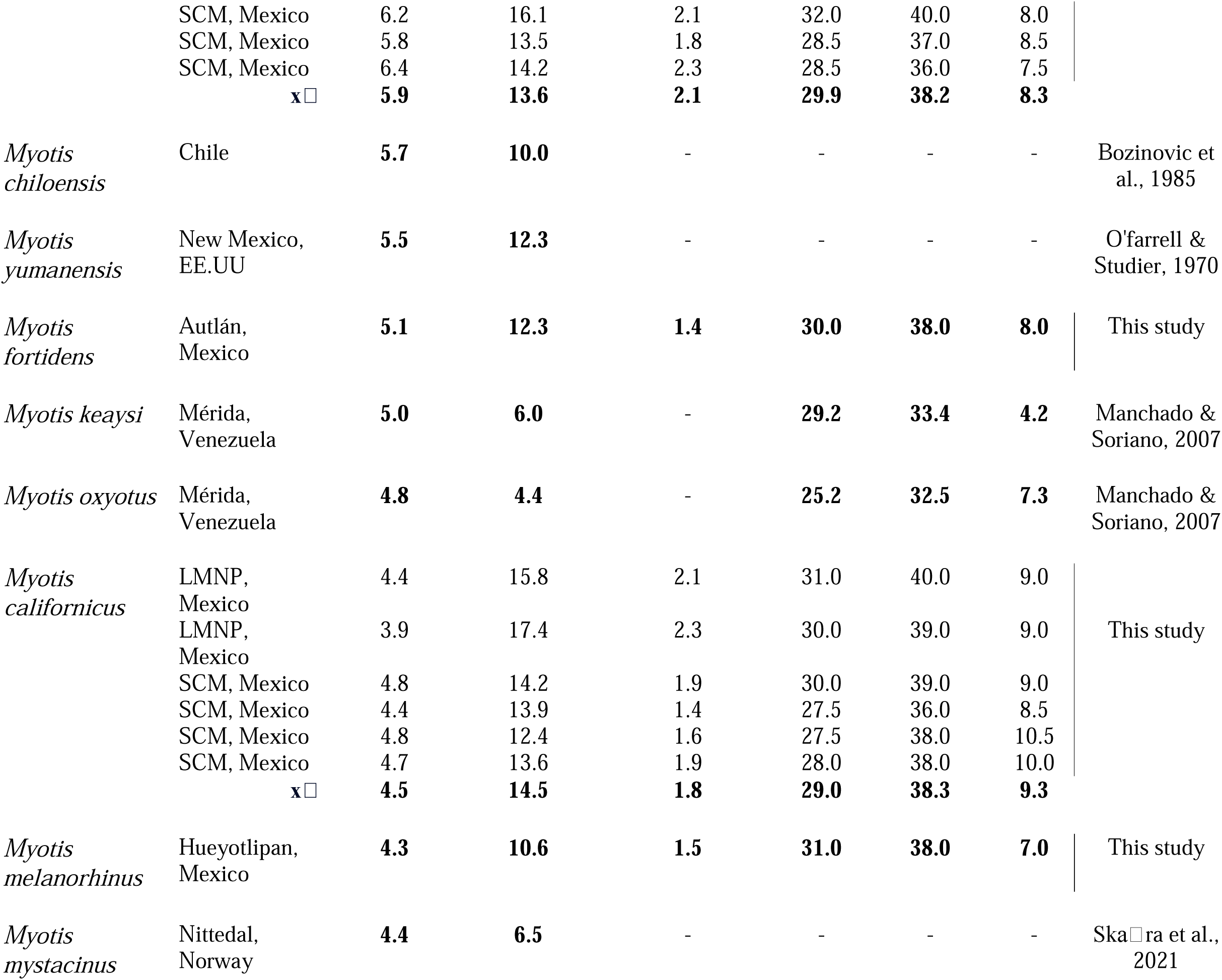
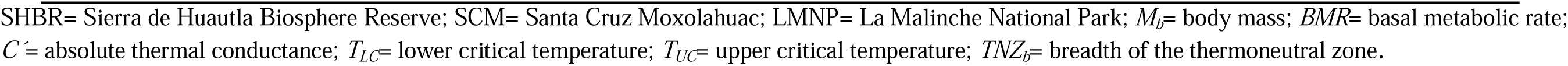
Thermal traits of the bat species of the family Vespertilionidae obtained from our metabolic measurements and those reported in the bibliography. Data presented as bold values (either mean or single values) are those that were used for the phylogenetic analyses. Data of locality from the study sites from the bibliography are presented as they could be retrieved in the published papers. nr denotes the information that was not reported.

Once captured, bats were transferred to laboratory conditions near the study sites, where they were placed in individual flight cages (75 x 75 x 75 cm) in temperature-controlled rooms (at a *T_a_* near the *TNZ* of bats at ∼ 28 °C; RH > 50 %; 12/12 light/dark cycle). The cages, made of plastic mesh, allowed bats to hang upside down as in natural conditions. Water was provided *ad libitum* to individuals before and after the experiments by holding each bat by hand and offering the liquid with a syringe. Bats were captured and handled with permission of the Department of Wildlife Management granted to our institution (SEMARNAT 07019; SGPA/DGVS/00582/20), and the University of Tlaxcala ethics committee.

### Determination of the thermal energetics

We estimated *BMR*, *T_LC_*, *T_UC_*, *TNZ_b_*, and *C*’ from the bats captured at the study sites. These variables were determined by measuring the resting metabolic rate of animals (in mL O_2_ h^-1^) over a range of *T_a_*’s from 8 to 43 °C. Measurements were taken from individuals in a postabsorptive state ∼10-12 hours after capture, during the resting phase (i.e., from ∼ 08:00 to 19:30 hours) (Genoud et al., 2018). Keeping the bats for a few hours reduced the potential effects of captivity on thermal physiology (Geiser & Brigham, 2000). We estimated the bats’ O_2_ consumption and CO_2_ production using open-flow respirometry (FoxBoxⓇ, Sable Systems International, Las Vegas, NV, USA.). We used O_2_ consumption and CO_2_ production because they guaranteed the most accurate estimation of the individuals’ metabolic rate (McNab, 2012). Measurements at each *T_a_* were taken once the readings of the two gasses reached an asymptote, which was achieved once that the *T_a_* stabilized within the chamber (i.e., ± 0.5 °C around the mean value of each experimental *T_a_*). Measurements were done following Medina-Bello et al. (2023). To do this, we placed each bat in a metabolic chamber (410 mL) inside a cabinet connected to a digital temperature controller (PELT5®; Sable Systems International, Las Vegas, USA). The chamber had plastic mesh on the walls and ceiling to allow the bats to crawl and hang upside down as in natural conditions. However, due to the dimensions of the chamber, the bats were unable to fly. This allowed us to minimize the variation in the metabolic measurements due to the movement of individuals. The *T_a_* inside the cabinet was controlled within ± 0.5 °C. Flow rates of dry CO_2_-free air scrubbed with Drierite (calcium sulfate for moisture), and Ascarite (sodium hydroxide for CO_2_) passed upstream with the air pushed through the respirometry chamber at a rate between 200- and 300-mL min^-1^ depending on the bats’ metabolic rate. Flow rates (FR) were calculated following Lighton & Halsey (2011) using the formula:

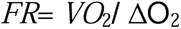

Where *VO*_2_ is the predicted oxygen consumption of the experimental animal and ΔO_2_ is the difference of the fractional concentration between incurrent and excurrent O_2_. *VO*_2_ was estimated assuming that bats scale their metabolic rate with *M_b_* with a slope of 0.744 (log_e_ *BMR* (mL O_2_ h^-1^) = 1.0895 + 0.744 log_e_ *M_b_* (g)) (Speakman & Thomas, 2003). ΔO_2_ was taken from our FoxBox analyzer (0.05 % or 0.0005 of ΔO_2_ expressed as fractional concentration). Excurrent air was dried with Drierite, and fractional concentrations (%/100) of both oxygen (*F_e_0*_2_) and CO_2_ (*F_e_C0*_2_) were measured every second. We placed one empty chamber of the same size (410 mL) inside the cabinet to take simultaneous baseline measurements from the animal chamber every second. Because we did not have any system that allowed us to switch the airflow between the empty chamber and the animal’s chamber to the same respirometer, we followed the protocol used by Medina-Bello et al. (2023) to take the baseline measurements in our experiments. In this methodology, the empty chamber was connected to a different respirometer of the same brand (FoxBox®, Sable Systems International, Las Vegas Nevada, U.S.A.) with the same type of sensors (O_2_ fuel cell and CO_2_ infra-red sensors) and the same flow rate as that of the experimental chamber. The two respirometers were calibrated simultaneously and showed the same pattern of oxygen readings (regression formulas obtained for the relationship between the O_2_ readings and the *T_a_*: O_2_ reading = −0.0028 *T_a_* + 21.23 and O_2_ reading = −0.0027 *T_a_* + 21.22 for the baseline respirometer and the animal respirometer, respectively) when the respirometers were tested with empty chambers at the same *T_a_*’s used in our experimental design (8, 13, 18, 23, 28, 33, 38 and 43 °C, for 30 min at each *T_a_*). This allowed us to correct our metabolic measurements from the drift using the data obtained from the baseline chamber.

We placed each bat inside the chamber at a *T_a_* of 28 °C ∼ 60 min before taking any measurement. The first *T_a_* tested for each individual was 28 °C, which we have previously determined falls within the *TNZ* of all bat species. We measured the metabolic rate of bats for 30 min and decreased the *T_a_* to 8 °C, taking measurements for 5 min for each drop in 1 °C and 30 min at each 5 °C (i.e., at 23, 18, 13, and 8 °C). Then, we raised the *T_a_*of the chamber to 28 °C for 30 min before increasing the *T_a_*to 43 °C, taking measurements for 5 min for each increase in 1 °C, and 30 min at each 5 °C (i.e., at 33, 38, and 43 °C). It took 5 to 10 min for the *T_a_* to stabilize for each change in *T_a_* within the metabolic chamber. The metabolic measurements we obtained from the animals at each °C allowed us to obtain a more precise estimation (1 °C rather than a 5 °C accuracy) of the *BMR*, *C*’, *T_LC_ T_UC_*, and *TNZ_b_*. One bat was measured at each time. We increased the *T_a_* inside the metabolic chamber and repeated the measurements if the bats reduced their metabolic rate below their *BMR* and intended to use torpor. With this methodology, bats from all the sites increased their resting metabolic rate when the *T_a_* fell below their *TNZ*, indicating that they defended normothermia. The *T_a_* within the metabolic chamber and the empty chamber were recorded using thermocouples connected to the FoxBox devices. Data from the FoxBox, which included the O_2_ and CO_2_ readings, as well as the flowmeter and thermocouple readings, were sent to computers through Sable Systems-UI2 ports running the Sable Systems Expedata software. After completing the measurements, we released each bat in the evening at their capture site.

#### Data analyses

All analyzes were performed in the R software version 4.1.0 (R Core Team, 2021). We measured the metabolic rate of bats through oxygen consumption (*VO*_2_) (in mL O_2_ h^-1^) for each bat tested at each *T_a_*. The metabolic rate was calculated by using the formula:

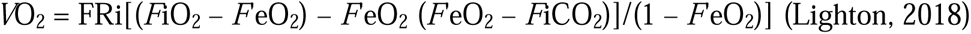

Where *FR* is the flow rate (in mL min^-1^), *F_i_ 0*_2_ is the fractional concentration of *0*_2_ in the incurrent (baseline) air, *F_e_0*_2_ is the fractional concentration of *0*_2_ in the excurrent air, *F_l_ C0*_2_ is the fractional concentration of *C0*_2_in the incurrent air, and *F_e_C0*_2_ is the fractional concentration of *C0*_2_ in the excurrent air. To calculate thermal variables, we used the mean values of the metabolic rate obtained from the 30 min measurements of the experimental *T_a_*’s and the mean values of the 5 min obtained from the animals at each °C. We defined the *BMR* (in mL O_2_ h^-1^) as the metabolic rate experienced by the bats between *T_LC_* (°C) and *T_UC_* (°C) (Genoud et. al., 2018, Geiser, 2021). Critical temperatures were estimated by continuous iterative two-phase regressions using the “chngptm” function (“chngpt” package) (Fong et al., 2017). These models estimate the points or thresholds of abrupt changes in the relationship between the dependent and independent variables (Nickerson et al., 1989). In bats, two-phase regressions have been used to calculate thermal traits, such as the *BMR*, *T_LC_*, and *T_UC_* (see Willis et al., 2005a; 2005b; Machado & Soriano, 2007, among others). In our models, the metabolic rate of bats was the dependent variable, and *T_a_* was the independent one. Finally, absolute *C*’ (in mL O_2_ h^-1^ °C^-1^) was calculated as the slope of the relationship between metabolic rate and *T_a_* below the *T_LC_*(McNab, 1980; Speakman & Thomas, 2003).

To conduct our analyses, we used the mean values of the metabolic rate we obtained from our experimental measurements and the data of the thermal variables we gathered from the bibliography for other members of the family Vespertilionidae (Table 2). To do this, we performed a systematic search in google scholar and the web of knowledge using the following keywords: “bat basal metabolic rate”, “bat lower and upper critical temperature”, “bat thermal energy”, “bat metabolism”, “bat thermal conductance”, “bat resting metabolic rate”, and “thermal physiology of bats”. We only included data of the thermal traits of those investigations that have used a similar methodology of that we performed in this work (i.e., with the use of open flow respirometry of non-reproductive postabsorptive adult animals measured over a range of *T_a_*’s). As a result, we obtained different samples to be used for the analyses for the different thermal variables: *BMR* (*n* = 38), *T_LC_* (*n* = 21), *T_UC_* (*n* = 18), *ZTN_b_* (*n* = 19) (Table 2). Unfortunately, we did not find any useful information for the *C*’ of any bat species as all results reported by other authors were calculated using the body temperature of the individuals. So, for this thermal variable, we only used our experimental data (*n* = 15).

#### Comparative analysis

To make our results comparable with previous studies, all analyses were conducted in the log_e_ transformed data. We evaluated the relationship of *M_b_*with the thermal traits using phylogenetic generalized least squares models (PGLS) using the function “gls” (package “nlme” version 3.1-153) (Pinheiro et al., 2021). The PGLS is a regression model that fits the covariance matrix considering the non-independence of the data due to the species evolutionary history (Felsenstein, 1985; Paradis, 2006). Besides, it has been observed that phylogenetic analyses to evaluate the relationship between thermal variables and *M_b_* fit the data better than conventional linear regressions (Dann et al., 1990; Riek & Geiser, 2013; Ayala-Berdon & Medina-Bello, unpublished data; Jetz et al., 2008). In the models, the thermal variables were the dependent variables and the *M_b_*the independent one. The regressions were conducted assuming a Brownian evolutionary model (BM) (Münkemüller et al., 2012). We calculated the phylogenetic signal index “λ” (corresponding to the BM), which is determined by the maximum likelihood “ML” estimate from the models (Pagel, 1999; Freckleton et al., 2002). This index ranges from 0 to 1, where values close to 1 indicate a strong phylogenetic signal and values close to 0 indicate a lack of phylogenetic signal in the trait (Freckleton et al., 2002). In our comparative analysis, we used the phylogenetic tree published by Shi & Rabosky (2015), which is composed of more than 800 species of bats from the different families of the order Chiroptera. Because we were interested only in the species of the family Vespertilionidae, we cut the tree using the “picante” package (Kembel et al., 2010). To evaluate the models’ fit, we extracted the AIC values from the summaries from the models.

#### Constructing the thermal predictive models

In this work we found that *BMR*, *C*, *T_LC_*, and *T_UC_* of bats were related to *M_b_* in our models (see the results section). So, we generated predictions of the allometric relationships of thermal traits in the phylogeny for the members of the family Vespertilionidae (including species of the genera *Cistugo* and *Miniopterus*) and Molossidae (see below). We generated the values of the thermal traits using the intercept and slope we obtained from the PGLS models and the average *M_b_* data reported for the 407 bat species available in the panTHERIA (Jones et al., 2009), Elton Trait (Wilman et al., 2014) and Moyers Arévalo et al. (2018). As the taxonomic identity of some bat species has changed over time, we used the *M_b_* value using the arrangement initially proposed by the Elton Trait database. We trimmed the phylogenetic tree published by Shi & Rabosky (2015) using the “picante” package (Kembel et al., 2010). The resulting phylogenetic tree had 239 bat species, as we did not obtain phylogenetic information for the rest of the species. We also obtained data from three species of the family Molossidae belonging to the genus *Tadarida* to be used as outgroups. Finally, we explored the evolution of the thermal traits by reconstructing the ancestral states and calculating the variance and confidence intervals (CI’s) using the “fastANC” maximum likelihood algorithm (Revell, 2012) with the function “anc.recon” (package “Rphylopars”; Goolsby et al., 2021) and mapped them on the phylogeny of the family Vespertilionidae using the contmap function from Phytools (Revell, 2012).

## RESULTS

### Thermal traits and M_b_

In this section, data from the phylogenetic analyses are presented as single values, while data from the extant bat species are presented as means with their respective standard error. We found that most of the thermal traits were related to *M_b_*, and the models showed medium-to-relatively-high phylogenetic signals. First, we found a positive relationship between *M_b_* and *BMR* (ML λ= 0.51, log_e_ *BMR* = 0.52 log_e_ *M_b_* + 1.65, t= 4.15, df= 36, p < 0.001, AIC: 44.09) (Fig. 2a). In this relationship, the smaller *M_b_* species *Myotis mystacinus* (4.4 g) and *M. oxyotus* (4.8 g) presented low values of *BMR* (6.5 and 4.4 mL O_2_ h^-1^, for *M. mystacinus* and *M. oxyotus*, respectively) while the larger *M_b_*species *Eptesicus serotinus* (27.0 g) and *Nyctalus leisleri* (30.3 g) showed higher values of this thermal trait (43.1 and 30.3 mL O_2_ h^-1^, for *E. serotinus* and *N. leisleri*, respectively). Nevertheless, we found that the *BMR* values of the smaller *M_b_ E. furinalis*, *Rhogeessa parvula* and *R. alleni* were very high (3.8, 5.5 and 5.9 times respectively higher than those expected from their *M_b_*) (Table 2). When we removed those species from the analysis, we found a better fit for the model (AIC: 26.23), a higher phylogenetic signal (ML λ= 0.71), and the estimates (log_e_ *BMR* = 0.72 log_e_ *M_b_* + 1.10) were almost identical of those published by Speakman & Thomas (2003) for a mix of bats belonging to 11 families of the order Chiroptera (log_e_ *BMR* = 0.74 log_e_ *M_b_* + 1.08), and similar of what Skåra et al. (2021) reported for bats of the family Vespertilionidae (log_e_ *BMR* = 0.71 log_e_ *M_b_* + 0.37). Therefore, we used the estimates of the model that excluded *E. furinalis*, *R. parvula* and *R. alleni* to construct the predictions of the allometric relationship between *M_b_* and the *BMR* and the ancestral reconstruction for the *BMR*.

**FIGURE 2.**
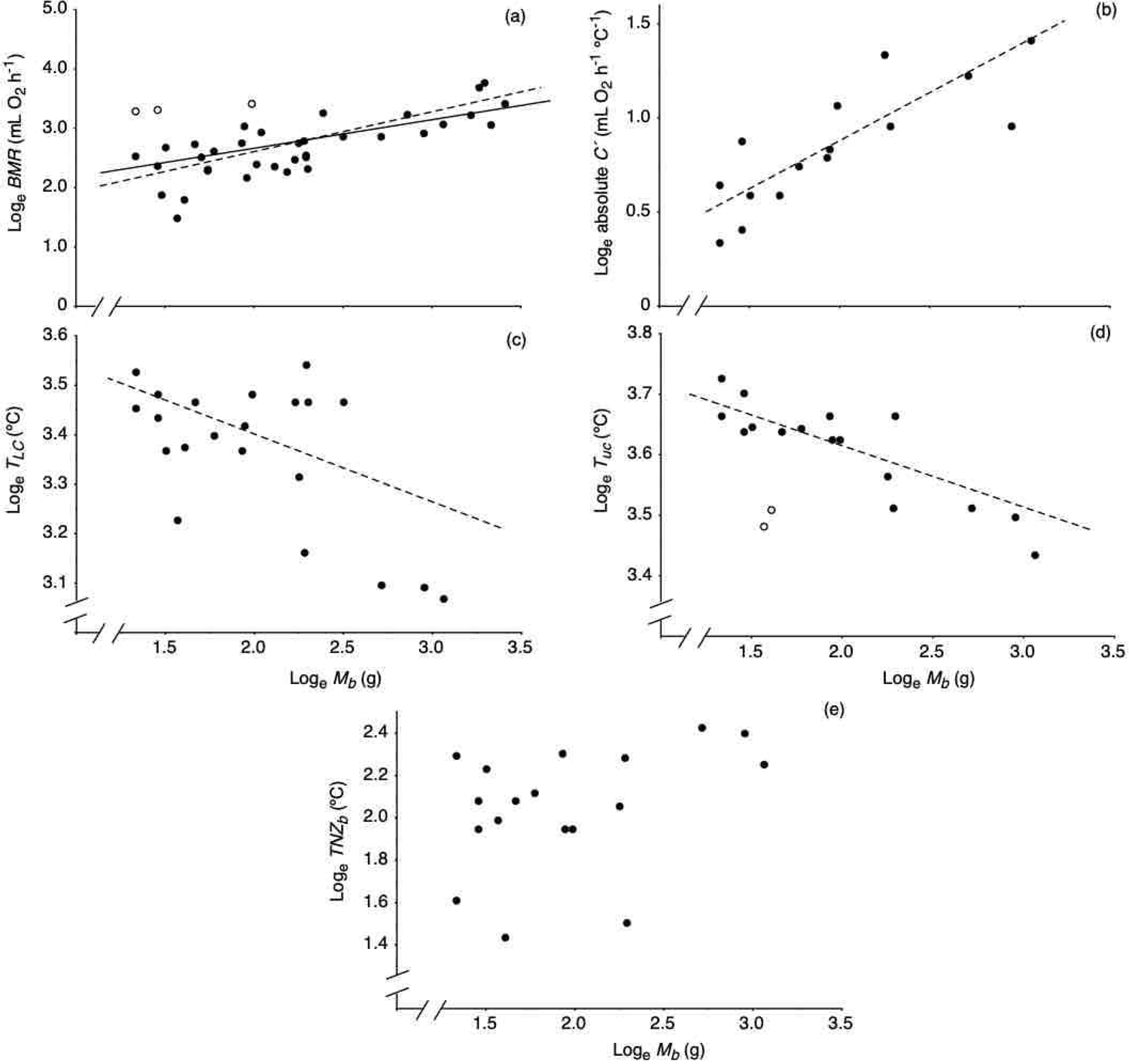
(a) Relationship between body mass (*M_b_*) and basal metabolic rate (*BMR*), (b) absolute thermal conductance (*C’*), (c) lower critical temperature (*T_LC_*), (d) upper critical temperature (*T_UC_*), and (e) breadth of the thermoneutral zone (*TNZ_b_*) of bats of the family Vespertilionidae. Dots represent the mean or single values of the data we obtained from our experimental trials and data we gathered from the bibliography. The trend lines were calculated using the parameters we obtained from the phylogenetic models. For the *BMR* regression, the solid line represents the relationship of the variables that included *Rhogeessa alleni*, *R. parvula*, and *Eptesicus furinalis* (represented by open dots), that presented very high values of *BMR* of those expected by their *M_b_*, while the dashed line denotes the regression that excluded these species. For the *T_UC_* regression, open dots depict *Myotis oxyotus* and *M*. *keaysi* which presented very low values of this thermal trait.

We also found a positive relationship between *M_b_* and absolute *C*’ (ML λ= 0.69, log_e_ *C*’ = 0.48 log_e_ *M_b_* −0.09, t= 3.56, df= 13, p= 0.003, AIC: 2.41) (Fig. 2b). In this relationship, the smaller *M_b_*species *R. tumida* (3.8 g) and *R. alleni* (3.8 g) showed low values of *C*’ (1.4 and 1.9 mL O_2_ h^-1^ °C^-1^, for *R. tumida* and *R. alleni*, respectively) while the larger *M_b_* species *E. fuscus* (15.1 g) and *Lasiurus cinereus* (21.4 g) presented high values of this thermal trait (3.4 and 4.1 mL O_2_ h^-1^ °C for *E. fuscus* and *L. cinereus*, respectively). The *T_LC_*of bats was negatively related to *M_b_* (ML λ= 0.62, log_e_ *T_LC_*= −0.17 log_e_ *M_b_* + 3.73, r^2^ = 0.25, t= −2.29, df=19, p = 0.03, AIC: −19.82) (Fig. 2c). In this relationship, the smaller *M_b_* species *R. alleni* (3.8 g) and *R. parvula* (4.3 g) presented values of *T_LC_* at warmer *T_a_’s* (31.6 and 34.0 °C for *R. alleni* and *R. parvula*, respectively) while the larger *M_b_* species *L. intermedius* (19.2 g) and *L. cinereus* (21.4 g) showed *T_LC_* values at colder *T_a_’s* (22.0 and 21.5 °C for *L. intermedius* and *L. cinereus*, respectively). We found a marginal non-significant relationship between *M_b_* and the *T_UC_*of bats (t= −1.98, df= 16, p=0.06, AIC: −44.50) (Fig. 2d). However, we identified very low *T_UC_* values for the small *M_b_* species *M. oxyotus* and *M*. *keaysi* which acted as influential points in our analysis (Fig. 2d). When we removed these values, we found a negative relationship between *M_b_* and the *T_UC_*of individuals (ML λ= 0.57, log_e_ *T_UC_*= −0.11 log_e_ *M_b_* + 3.83, r^2^ = 0.81, t= - 4.94, df=16, p < 0.001, AIC: −54.75) (Fig. 2d). In this relationship, the smaller *M_b_* species *R. alleni* and *R. parvula* presented values of *T_UC_* at warmer *T_a_’s* (41.5 and 40.5 °C for *R. alleni* and *R. parvula*, respectively) while the larger *M_b_* species *L. intermedius* and *L. cinereus* showed *T_UC_* values at colder *T_a_’s* (33.0 and 31.0 °C for *L. intermedius* and *L. cinereus*, respectively). Because it has been found that *M. oxyotus* and *M*. *keaysi* have displaced their *T_UC_* towards colder *T_a_*’s as an adaptation to live in cold environments in highlands in the Venezuelan Andes (Machado & Soriano, 2007), we used the estimates of the model that excluded those species to construct the predictions of the allometric relationship between *M_b_* and *T_UC_* and the ancestral reconstruction for the *T_UC_* of bats of the family Vespertilionidae. Finally, we did not find any relationship between *M_b_* and *TNZ_b_* of bats (t= 1.59, df= 16, p=0.12, AIC: 15.57) (Fig. 2e).

### Ancestral state reconstruction of the thermal variables and M_b_

The state reconstruction of the *M_b_* for the common ancestor of bats of the family Vespertilionidae showed an intermediate value of 15.0 g (see supplementary material II). However, we observed a lower mean value of 10.4 ± 0.4 g for extant bat species and identified similar values to those of the ancestral estimate in some species like *Pipistrellus stenopterus*, *Myotis welwitschii*, *Euderma maculatum* and *Barbastella leucomelas*. *Tadarida fulminans*, *Scotophilus heathii*, *Nyctalus lasiopterus* (forest, temperate), *Myotis chinensis* and *La io* independently evolved higher *M_b_* (> 30.0 g), while some species of the genera *Kerivoula*, *Myotis*, *Neoromicia*, and the species *Murina suilla* and *Tylonycteris pachypus* independently evolved lower *M_b_* (< 4.0 g) (see supplementary material II).

Because our predicting models were constructed based on the relationship between the thermal energetics and *M_b_*, we found the same pattern of the evolutionary changes in our thermal variables. The ancestral estimate of *BMR* and *C*’ presented intermediate values of 20.6 mL O_2_ mL h^-1^ and 3.2 O_2_ mL h^-1^ °C^-1^, respectively. However, the average values for extant bat species were lower (15.7 ± 0.3 mL O_2_ h^-1^ and 2.7 ± 0.0 mL O_2_ h^-1^ °C^-1^ for the *BMR* and *C*’, respectively). The ancestral estimate for *T_LC_*showed an intermediate value of 27.1 °C, while the average value for extant bat species was 28.8 ± 0.1 °C (see supplementary material I). The ancestral estimate for *T_UC_*showed an intermediate value of 35.1 °C, while the average value for extant bat species was 36.5 ± 0.1 °C (see supplementary material I). In our models, we observed changes of *BMR* from the ancestral estimate to values ≥ 39.6 O_2_ mL h^-1^ and ≥ 5.1 mL O_2_ h^-1^°C^-1^ in *C*’ in extant bat species > 30 g, and changes on *BMR* to values ≤ 8.0 mL O_2_ h^-1^ and ≤ 3.6 mL O_2_ h^-1^ °C in *C*’ in extant bat species < 4 g. We also found changes from the ancestral estimate on *T_LC_*and *T_UC_* towards lower *T_a_*’s ≤ 22.4 °C and ≤ 32 °C respectively in extant bat species > 30 g, and changes to higher *T_a_*’s ≥ 32.8 °C and 41.5 °C respectively in extant bat species < 4 g.

## DISCUSSION

### Thermal energetics and M_b_

In this work we found that most thermal variables were related to *M_b_* in our study species. First, we found a positive relationship between *M_b_* and *BMR*. This is similar to what has been found for 639 species of mammals including bats of different families (McNab, 2008). Since the first description made by Rubner (1883), the allometric effect of *BMR* has been explained by simple physical and geometric equations. Because the heat produced by an animal is lost to the environment through the body surface, it has been proposed that the differences on *BMR* are proportional to differences in the area-to-volume ratio of the animal species (which should be *M_b_* ^0.66^). However, many studies have shown that *BMR* on a variety of mammals escalates at a rate ∼ *M_b_* ^0.75^ (Kleiber, 1932; Brody, 1945; Savage et al., 2004; Kozłowski et al., 2003; Farrell-Gray & Gotelli, 2005, among others). This can be explained because not all mammals have a sphere-shape in their bodies, as the area-to-volume ratio hypothesis assumes (Mould, 2006; West et al., 2015; Riemer et al., 2018). At the beginning of the XXI century, Brown et al. (2003) proposed a very interesting theory on the basis of the fractal arrangement of the organisms. According to the authors, the way in which the nutrients and the materials necessary for the maintenance of the animals’ life are distributed in their own body (e.g., by the circulatory and respiratory systems which are structured as fractals) can explain how the *BMR* scales with *M_b_* ^0.75^. Nevertheless, because of some flaws in its theoretical foundations and the fact that not all organisms are composed completely by fractals, many criticisms have been raised to this theory (Kozłowski & Konarzewski, 2004; Makarieva et al., 2005; Apol et al., 2008). Additionally, many other studies have found that the *BMR* of certain groups of animals escalate with different slopes (i.e., from 0.66 to 1) (Hayssen & Lacy, 1985; Dodds et al., 2001; Symonds & Elgar, 2002; White & Seymour, 2003, 2005; Glazier, 2005, 2006; Painter et al., 2005; White et al., 2006; Duncan et al., 2007; White et al., 2007; Packard & Birchard, 2008; McNab, 2008; Sieg et al., 2009; White et al., 2009). In our models, we found a lower exponent of the relationship between *M_b_* and *BMR* (*M_b_*^0.51^) than those reported in the past. This can be explained by the surprisingly high values of *BMR* we found for *R. alleni*, *R. parvula* and *E. furinalis*, which acted as influential points in our regression analysis. When we excluded these species from the model, we found a better fit and an exponent of *M_b_* ^0.71^ which is very similar to other values published for bats in the past (e.g., Speakman & Thomas, 2003; Skara et al., 2021). Nevertheless, the very high *BMR* values we found for *R. alleni*, *R. parvula* and *E. furinalis* are quite interesting because, to the best of our knowledge, such high values had never been reported for bats of any family of the order Chiroptera. This can be the result of the research on thermal energetics biased to bat species from temperate regions of the word. So, more studies from the tropics and subtropics on this topic need to be conducted. Although it has been shown that the *BMR* of insectivorous bats tends to be lower than bats having other diets like frugivory or nectarivory (Speakman & Thomas, 2003), the high values of *BMR* presented here for *R. alleni*, *R. parvula* and *E. furinalis* are only comparable to those shown from small shrews of the family Soricinae (with *M_b_* ranging from 3.3 to 7.7 g), which have been reported to present the highest *BMR*’s among mammals, with values from three to six times higher than expected from their *M_b_* (Morrison et al., 1959; Sparti and Genoud, 1989; Nagel, 1994; Taylor, 1998). In shrews, elevated values of *BMR* have been related with high levels of thyroxine (Tomasi, 1983) a hormone that has been identified affecting the metabolic rate of other mammals and birds (Banta & Holcombe, 2002). Nevertheless, there is no information of the relationship between the thyroxine levels and *BMR* in bats. Although the levels of thyroxine or other hormones may explain the high values of *BMR* found in some of our study species, this topic needs further exploration.

In this investigation, we found that *M_b_* of bats was positively related with absolute *C*’. This is similar to what has been reported for 86 species of mammals and 93 species of birds in other studies conducted in the past (see Aschoff, 1981 and the references cited there). Interestingly, the exponent of the relationship we found (*M_b_* ^0.49^) was almost identical of those derived by Herreid & Kessel (1966) (*M_b_* ^0.51^), Aschoff (1981) (*M_b_* ^0.51^) and McNab (2002) (*M_b_*^0.50^) for other mammal species. Differences in *C*’ among bats varying in *M_b_* can be explained by the way that heat is transferred from the individuals’ core to the environment (Speakman & Thomas, 2003; McNab, 2021). In animals, heat flow is transferred internally by convection via blood flow through the circulatory system, by conduction through the cells and tissues, and by convection, conduction, radiation, and evaporation through the skin surface (McNab, 2002). All those components (including the fur attached to the skin) act as resistance layers to heat flow, which is the inverse of *C*’. This means that larger species should present higher resistances and therefore lower *C’*, which is the opposite of what we found for our study species. This is explained simply because we measured absolute *C*’, which is higher in large animals just because of their larger size. Nevertheless, as we discussed before, the way that heat is transferred to the environment is also dependent on *M_b_*, because heat loss occurs over the body and surface which is larger in small animals. This results in higher *C’* values corrected per *M_b_* in small mammals (i.e., the so-called mass specific *C*’). In our study species, larger bats present more resistant layers per gram because have larger circulatory systems, organs, muscles, and a skin surface-area that can act as resistance paths to heat flow (i.e., by having lower specific *C*’) while small bats that have short path lengths between the core and skin surface per unit of area, present high rates of heat flow because of their high mass specific *C*’ (Herreid & Kessel, 1967; Aschoff, 1981).

Here we found a negative relationship between *T_LC_* and *T_UC_* and the *M_b_*of bats. This result is similar to what Riek & Geiser (2013) reported for both critical temperatures of 204 mammal species compiled from the bibliography. In their phylogenetic analysis, the authors found an exponent of the relationship between *T_LC_*and *M_b_* (0.20) for placentals that was similar to the value we obtained here (0.17). Nevertheless, the exponent for the *T_UC_* we obtained (0.11) was larger than that reported for placentals by the authors (0.03). This suggests that, for bats of the family Vespertilionidae, the *T_UC_* escalates higher than for the rest of placentals. The negative relationship between *M_b_* and critical temperatures can be explained by: 1) the interaction of *BMR* and *C*’ that affect the *T_LC_* (Speakman & Thomas, 2003), and 2) the fur thickness (which affects the heat dissipation) that has an effect on the *T_UC_* of animals of different *M_b_*(Reik & Geiser, 2013). For example, it has been reported that smaller *M_b_* individuals with high *BMR* and *C*’ (which are also *M_b_* dependent) tend to present warmer *T_LC_*’s than larger ones (Bradley & Deavers, 1980; Capellini et al., 2010). Additionally, larger animals tend to present ticker fur that allows them to lose less heat with respect to the environment. However, dense fur in large animals may prevent heat dissipation which is highly related to *T_UC_*(Riek & Geiser, 2013). Differences in critical temperatures may have important implications for the geographic distribution and energy-saving strategies used by the animals, as these thermal traits define the *T_a_*’s at which individuals start expending energy for thermoregulation (Scholander et al., 1950; Fristoe et al., 2015; Mitchell et al., 2018). These implications will be discussed in an evolutionary context in the last part of the manuscript.

Finally, in this work we did not find any relationship between *M_b_* and *TNZ_b_* of bats. This result can be explained because, in our data, we found many bat species of similar sizes presenting a high variety of *TNZ_b_*. Differences in *TNZ_b_* can be more related to the climate where animals live. For instance, Bozinovic et al. (2014) found that, for 85 species of rodents, the species inhabiting cold climates presented a wider *TNZ_b_*than those living in warm climates. Similarly, Medina-Bello et al. (2023) found that individuals of the cave myotis (*M. velifer*) from a warm tropical dry forest had a narrower *TNZ_b_*than bats from a temperate forest. Although differences in climate affinity may explain the lack of relationship between *M_b_*and *TNZ_b_*, this topic needs further investigation.

### Thermal ancestral state reconstruction and M_b_

The most recent evolutionary hypotheses have estimated that bats have experienced several changes in *M_b_* once individuals started to fly (Hutcheon & Garland, 2004; Safi et al., 2005; Giannini et al., 2012; Moyers Arevalo et al., 2018). According to these hypotheses:1) bats experienced a generalized *M_b_* reduction while refining their capacity to fly. Likewise, here we found that the estimated common ancestor of bats of the family Vespertilionidae had a *M_b_* of 15 g and the extant bat species presented a mean *M_b_* value of 10.4 g. These values are comparable to those reported by Cabrera-Campos et al. (2021) for bats of the family Vespertilionidae (15 g and 9.6 g for the common ancestor and the extant bat species, respectively). 2) The reduction in *M_b_* was followed by a stasis among a median value of *M_b_*, and subsequent decreases and increases in *M_b_* in specific clades of the phylogeny (Hutcheon & Garland, 2004; Safi et al., 2005; Giannini et al., 2012; Moyers Arévalo et al., 2018). Similarly, in this work we identified five species that independently evolved *M_b_* > 30 g, and three genera and two species that evolved *M_b_* < 4.0 g. These differences may account for the high diversity of *M_b_* that can be found within bats of the family Vespertilionidae, which may vary as much as one order of magnitude among species (Jones & Purvis, 1997; Safi et al., 2013). The changes in *M_b_* throughout the evolutionary history of bats may have affected many of their life-history traits (Moyers Arévalo et al., 2018; Cabrera-Campos et al., 2021). Because there is strong evidence that *M_b_* and some thermal traits as *BMR* have not evolved independently in mammals (White et al., 2019), the changes in *M_b_* along their evolutionary history should have resulted in modifications in thermal energetics of bats of the family Vespertilionidae.

The decrease in *M_b_* once bats began to fly should have resulted in a decrease of absolute *BMR* and *C*’. According to this hypothesis, here we found that the mean values of *BMR* and *C*’ were 23.7 % and 15.6 % lower than the estimates of their common ancestor. These changes would have affected the *T_LC_* of individuals. In this work, we found that the *T_LC_* of bats was 6.2 % higher than the estimate of their common ancestor. Nevertheless, small *M_b_*should have represented a thermoregulatory challenge for individuals because heat loss to the environment increases with increasing the area-to-volume ratio (Austad & Fischer, 1991). So, for most bats of the family Vespertilionidae, the reduction in *M_b_* should have increased their heat loss with respect to the ecosystems (Speakman, 2005). In this condition, animals should have been forced to increase their daily energy intake (Speakman & Thomas, 2003). However, a *M_b_* decrement could have reduced the animals’ capacity to acquire energy, as the intestinal length and nominal-surface-area tend to decrease proportionally with *M_b_* in bats (Klite, 1965; Caviedes-Vidal et al., 2007). To resolve this difficulty, it has been hypothesized that bats of the family Vespertilionidae have increased the apparent dry matter digestibility in their small intestines in response to the reduction of *M_b_* over their evolutionary history (Cabrera-Campos et al., 2021). Nevertheless, although with many differences among ecosystems, food availability tends to be highly variable with important energy bottlenecks in certain seasons of the year (Nagy et al., 1999). In these seasons, the use of energy-saving strategies such as torpor and hibernation, which consist in the reduction of the metabolic rate and *T_b_* to decrease the energy expenditures of animals, may have played a fundamental role in their survival (Geiser and Brigham, 2012; Nowack et al., 2017).

Finally, differences in *T_LC_* and *T_UC_*that are affected by *M_b_* could have influenced the geographic distribution and the use of energy-saving strategies of bats of the family Vespertilionidae. For instance, Riek & Geiser (2013) and Bozinovic et al. (2014) proposed that differences in *T_LC_* and *T_UC_* can explain partially the dominance of small species in warmer climates and larger species in colder climates among mammals and rodents. This trend seems to be similar in bats. For example, Alston et al. (2023) collected data from more than 30,000 individuals belonging to 20 species of bats (19 of them belonging to the family Vespertilionidae) over a decade in North America and found smaller *M_b_*bats in warmer climates than colder ones. Similarly, in our study sites bats from warmer environments presented *M_b_* values averaging 6.6 g compared to the 10 g of bats from the cold climates. Additionally, Ayala-Berdon & Medina-Bello (unpublished data) found that *T_LC_* was closely related to the *T_a_*at which 11 bat species of the family Vespertilionidae went into torpor when animals were energetically restricted. The authors also found smaller *M_b_* bats inhabiting warmer climates were able to use torpor at warmer *T_a_*’s. Although differences in *T_LC_* and *T_UC_* related to *M_b_* may determine the dominance of vespertilionid bats in the ecosystems and their use of energy-saving strategies, this topic calls for further exploration.

---

**FIGURE 3.** Ancestral reconstruction of the *M_b_* in the phylogenetic tree of the family Vespertilionidae. Red and yellow colors indicate low *M_b_* values, while blue and purple colors indicate high *M_b_* values.

**FIGURE 4.** Ancestral reconstruction of the *BMR* in the phylogenetic tree of the family Vespertilionidae. Red and yellow colors indicate low *BMR* values, while blue and pink colors indicate high *BMR* values.

**FIGURE 5.** Ancestral reconstruction of absolute *Ć* in the phylogenetic tree of the family Vespertilionidae. Red and yellow colors indicate low *Ć* values, while blue and pink colors indicate high *Ć* values.

**FIGURE 6.** Ancestral reconstruction of the *T_LC_* in the phylogenetic tree of the family Vespertilionidae. Red and yellow colors indicate low *T_LC_* values, while blue and pink colors indicate high *T_LC_* values.

**FIGURE 7.** Ancestral reconstruction of the *T_UC_* in the phylogenetic tree of the family Vespertilionidae. Red and yellow colors indicate low *T_UC_* values, while blue and pink colors indicate high *T_UC_* values.

